# Plasmid-mediated macrolide resistance among rapidly growing mycobacteria in Japan

**DOI:** 10.1101/2025.09.17.676775

**Authors:** Takeshi Komine, Panuwat Sathianpitayakul, Nobuya Sakagami, Mitsunori Yoshida, Masato Suzuki, Yoshihiko Hoshino, Panan Ratthawongjirakul, Manabu Ato, Hanako Fukano

## Abstract

**Objectives:** The spread of a transmissible plasmid carrying the 23S rRNA methylase gene *erm*(55), which confers inducible macrolide resistance in rapidly growing mycobacteria (RGM), has raised significant clinical concerns. The aim of this study was to investigate the prevalence of *erm*(55)-carrying plasmids in clinically isolated RGM strains in Japan.

**Methods:** In total, 607 RGM clinical isolates, representing 32 species or complexes, collected between 2019 and 2023 in Japan were examined. To detect the presence of *erm*(55)-carrying plasmids, we conducted PCR screening, minimum inhibitory concentration testing for clarithromycin, and whole-plasmid genome sequencing. Comparative genomic analyses were performed to characterise the plasmids.

**Results:** Among the 607 RGM isolates, 0.8% (5/607) possessed the plasmid with the *erm*(55) gene and exhibited inducible macrolide resistance, with ratios of 100% (1/1) in *Mycobacterium murale*, 50% (3/6) in *M. obuense*, and 0.8% (1/125) in *M. chelonae*. The *erm*(55)-carrying plasmids ranged from 126,187 to 170,220 bp in size. Pairwise BLASTn comparisons of the *erm*(55)-carrying plasmids showed weighted percent identity values ranging from 99.5% to 99.9%, with query and subject coverage values ranging from 74.2% to 100%. All *erm*(55) sequences (813 bp) were identical and located within a horizontal gene transfer region.

**Conclusions:** This study confirmed the presence of macrolide-resistant RGMs related to the *erm*(55)-carrying plasmid in Japan, although the overall prevalence remains low. These findings emphasise the need to consider plasmid-mediated resistance when treating infections caused by the RGM species.

## Introduction

The incidence of nontuberculous mycobacterial (NTM) infection is increasing worldwide.^1^ In Japan, the incidence of pulmonary NTM infection was 19.2 per 100,000 people in 2017, exceeding that of tuberculosis.^2^ In Asia, the proportion of rapidly growing mycobacteria (RGM) infections (e.g., *Mycobacterium abscessus* and *M. chelonae*) is fairly high among the NTMs.^3,4^

Treatment of NTM infections typically involves multidrug therapy, including macrolide antibiotics such as clarithromycin or azithromycin.^5^ However, several mycobacterial species can exhibit acquired or inducible resistance to macrolides, which is associated with mutations in the 23S rRNA (*rrl*) gene or the 23S rRNA methylase (*erm*) gene, respectively. Acquired resistance-related mutations at positions 2058 and 2059 (*Escherichia coli* numbering) of *rrl* have been reported in *M. abscessus* and *M. chelonae*. The *erm* genes have been identified in various RGMs as follows: *erm*(41) in *M. abscessus*; *erm*(38) in *M. smegmatis* and *M. goodii*; *erm*(39) in *M. fortuitum, M. boenickei, M. houstonense, M. neworleansense*, and *M. porcinum*; and *erm*(40) in *M. mageritense* and *M. wolinskyi*.^6^

In 2023, a plasmid (pMchErm55) containing *erm*(55) was discovered in *M. chelonae*. This plasmid has also been found on other RGMs, such as *M. obuense* and *M. iranicum*, and may have the potential for horizontal gene transfer within RGMs,^7^ representing a clinical concern for the treatment of NTM infections. Herein, we aimed to investigate the prevalence of *erm*(55)-carrying plasmids in clinical RGM isolates in Japan.

## Materials and methods

### Bacterial isolates

This study investigated 607 RGM strains isolated from a clinical microbiology laboratory in Japan between 2019 and 2023 (Figure S1, Table S1). The isolates originated from pulmonary (65.6%), extrapulmonary (8.2%), and other specimens (1.2%), whereas the origin of 25.0% of the isolates was unknown. These isolates included 32 different species or complexes that were identified using the MALDI Biotyper system (Bruker, Germany). To refine species identification within several complexes, average nucleotide identity analysis was performed.

### PCR screening

RGM isolates were screened by PCR using the primer set *erm*(55)P-F-1 and *erm*(55)P -R-1, targeting the *erm*(55)^P^ gene as described by Brown-Elliott et al.^7^ The heat shock protein 65 gene was amplified using the primer sets Tb11 and Tb12^8^ as positive controls (Supplementary methods).

### MIC determination

Inducible phenotypic macrolide resistance was assessed in isolates that tested positive for *erm*(55) by PCR, as well as in the type strain of *M. chelonae* (JCM 6388), using clarithromycin in accordance with the Clinical and Laboratory Standards Institute protocol.^9^ The minimum inhibitory concentration (MIC) values were determined on days 3 and 14.

### Sequencing and plasmid assembly

We extracted genomic DNA from mycobacterial isolates using a previously reported method^10^ and obtained sequencing data using Rapid Barcoding Kit 24 V14 and the P2 Solo platform with the R10.4.1 flow cell (Oxford Nanopore Technologies, UK) (Table S2). Base-calling was performed using Dorado v0.9.0 in super accuracy mode (https://github.com/nanoporetech/dorado/). Reads with >*Q*20 and >5000 bp were then extracted using NanoFilt v2.8.0^11^ and assembled using Flye v2.9.3-b1797.^12^ The plasmid and chromosome genomes were annotated using DFAST (https://dfast.nig.ac.jp) and deposited in DDBJ (Table 1, Table S2). Annotated *rrl* sequences were also examined for point mutations at positions 2058 and 2059 (*E. coli* numbering). The *erm* genes, ranging from *erm*(37) to *erm*(55), were searched against the genomes using BLASTn v2.5.0.^13^

**Table 1.** Results of the screening tests in this study.

| Strain | Species | *erm*(55)<sup>p</sup><br>PCR<sup>a</sup> | Clarithromycin MIC<br>(mg/L) |  | Nucleotide at<br>*rrl* position<sup>b</sup> |  | *erm*(55)-carrying<br>plasmid assembly<sup>c</sup> | Accession No. for<br>plasmid |
| --- | --- | --- | --- | --- | --- | --- | --- | --- |
|  |  |  | 3-day | 14-day | 2058 | 2059 |  |  |
| JCM 6388<sup>d</sup> | *Mycobacterium chelonae* | - | 0.5 | 0.5 | A | A | NA | NA |
| SRL2021-127 | *Mycobacterium chelonae* | + | 2 | >16 | A | A | + | LC872744 |
| SRL2019-498 | *Mycobacterium obuense* | + | 2 | 16 | A | A | + | LC872745 |
| SRL2021-291 | *Mycobacterium obuense* | + | 0.5 | >16 | A | A | + | LC872746 |
| SRL2023-024 | *Mycobacterium obuense* | + | 1 | >16 | A | A | + | LC872747 |
| SRL2023-035 | *Mycobacterium murale* | + | 8 | >16 | A | A | + | LC872748 |

MIC, minimum inhibitory concentration; NA, not applicable
<sup>a</sup>+: The 23S rRNA methylase (*erm*(55)<sup>p</sup>) and heat shock protein 65 (*hsp65*) genes were amplified in the PCR screening tests
<sup>b</sup>23S rRNA (*rrl*) gene sequence position in *Escherichia coli* numbering
<sup>c</sup>+: *erm*(55)-carrying plasmid sequence was assembled in long-read sequencing analysis
<sup>d</sup>Control strain in the MIC test

### Comparative analysis of erm(55)-carrying plasmids

We performed pairwise BLASTn comparisons among the plasmids assembled in this study and previously reported plasmid pMchErm55 (CP118918.1). The *erm*(55)-carrying plasmids were compared and visualized using Proksee.^14^ In the reference genome, mobile genetic elements and horizontal gene transfer (HGT) regions were detected using mobileOG-db^15^ and alien_hunter v1.7.^16^

Plasmids were reannotated using Prokka v1.14.6 (https://github.com/tseemann/prokka) and pangenome analysis was performed using Roary v3.13.0 (https://github.com/sanger-pathogens/Roary). Phylogenetic analyses of plasmid sequences were also performed (Supplementary Methods).

## Results

### Prevalence of the erm(55)-carrying plasmid

In our screening tests, 0.8% (5/607) of the RGM isolates possessed the plasmid containing the *erm*(55) gene and showed inducible macrolide resistance, with ratios of 100% (1/1) in *M. murale*, 50% (3/6) in *M. obuense*, and 0.8% (1/125) in *M. chelonae* (Table S3). In the PCR screening test for the *erm*(55) gene, five isolates (*M. chelonae* SRL2021-127, *M. murale* SRL2023-035, and *M. obuense* SRL2019-498, SRL2021-291, and SRL2023-024) showed positive results (Table 1, Figure S1). In four isolates (*M. chelonae* SRL2021-127 and *M. obuense* SRL2019-498, SRL2021-291, and SRL2023-024), the MIC values changed from susceptible levels (3-day MIC, 0.5–2 mg/L) to resistant levels (14-day MIC, 16–>16 mg/L) (Table 1). *M. murale* SRL2023-035 showed resistance at day 3 (8 mg/L), with increases in MIC values observed at day 14 (>16 mg/L). Five plasmids with the *erm*(55) gene (pErm55Mc1, pErm55Mo1, pErm55Mo2, pErm55Mo3, and pErm55Mm1) were detected from *M. chelonae* SRL2021-127, *M. obuense* SRL2019-498, *M. obuense* SRL2021-291, *M. obuense* SRL2023-024, and *M. murale* SRL2023-035, respectively. No mutations were detected at positions 2058 or 2059 (Table 1, Figure S2).

### Comparison of erm(55)-carrying plasmids

The assembled plasmid length was 126,187–170,220 bp, with 119–165 coding DNA sequences (CDSs) (Table S2). Pairwise BLASTn comparisons of the *erm*(55)-carrying plasmids showed weighted percent identity values ranging from 99.5% to 99.9%, with query and subject coverage values ranging from 74.2% to 100% (Table S4). The *erm*(55)^P^ gene sequence (813 bp) was detected in the six plasmids, with 100% identity, and located within the HGT region (Figure 1). Plasmids pErm55Mo3, pErm55Mc1, and pErm55Mm1 lacked a region comprising nine CDSs, corresponding to nucleotide positions 1–3,848 bp and 130,065–137,526 bp of pMchErm55 (CP118918.1). This region included CDSs predicted to encode proteins involved in DNA recombination and mobility, such as a tyrosine-type recombinase/integrase and GIY-YIG nuclease family protein, and was inferred to represent part of the HGT region. Predicted insertion sequence (IS) elements, including members of the IS*200*/IS*605* and IS*110* families, were detected in all plasmids (Figure 1, Table S5). pErm55Mm1 harbored an IS*1380* family transposase. A putative ATP-binding protein related to IS*5376* was identified in pErm55Mo1 and pErm55Mo2. pErm55Mo3 carried IS*21* and IS*256* family transposases. The plasmids pErm55Mm1 and pErm55Mo2 harbored mercuric reductase (*merA*) and organomercurial lyase (*merB*), which are involved in mercury resistance.

**Figure 1.**
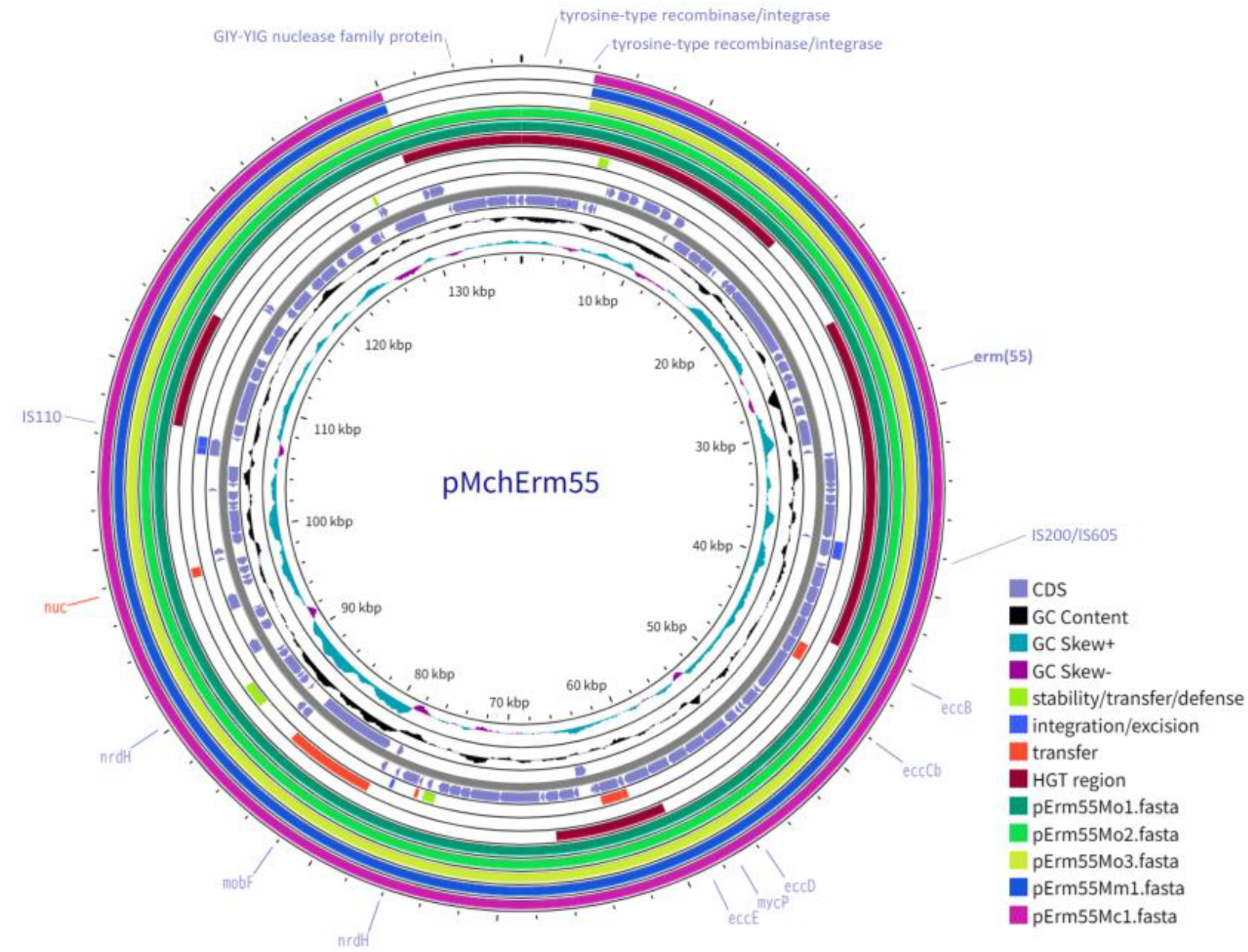
Comparative plasmid maps of *erm*(55)-carrying plasmids, the plasmids used in this study, pErm55Mo1, pErm55Mo2, pErm55Mo3, pErm55Mc1, and pErm55Mm1, and the previously reported plasmid, pMchErm55 (CP118918.1). The inner rings show the GC content and GC skew. Highlighted coding DNA sequences (CDS), plasmid functionality genes (stability/transfer/defense, integration/excision, transfer) from the mobileOG-db,^16^ and horizontal gene transfer (HGT) regions detected with alien_hunter v1.7.^17^ Created using ProkSee.com.^15^

Pangenome analysis detected 114 core genes among the six plasmids (Figure S3). In the core gene-based and recombination-free phylogeny (Figure S3), pMchErm55, identified from *M. chelonae*, was more closely related to pErm55Mm1 and pErm55Mo plasmids from other mycobacterial species than to pErm55Mc1, which was also derived from *M. chelonae*.

## Discussion

In this study, five isolates from three mycobacterial species, *M. chelonae, M. obuense*, and *M. murale*, showed inducible phenotypic macrolide resistance associated with *erm*(55)-carrying plasmids. At 0.8%, the proportion of *erm*(55)-positive isolates among *M. chelonae* clinical isolates in Japan from 2019 to 2023 was lower than that reported in the USA from 2019 to 2021 (3.8%).^8^ *M. obuense* showed a higher prevalence of the plasmid-associated *erm*(55) gene and *M. murale* can also harbor *erm*(55)-carrying plasmids. Although *M. obuense* and *M. murale* rarely cause human infections,^17,18^ the presence of the *erm* gene warrants consideration in therapeutic decision-making. Furthermore, the fact that these low-pathogenicity hosts carry such plasmids suggests the potential for horizontal transfer to more pathogenic species, including *M. chelonae*, which may have clinical implications.

This *erm*(55)-carrying plasmid suggests possible transmissibility across RGMs.^7^ Comparative analysis revealed plasmid diversity in terms of length and CDS content and showed that plasmids from different mycobacterial species were more closely related than those from the same species, supporting the possibility of interspecies transmission. Furthermore, the finding that these large plasmids harbor not only drug resistance genes but also heavy metal resistance genes implies a broad contribution to host environmental adaptability.

As suggested by Vial et al.,^19^ plasmid–chromosome interactions lead to adaptive genomic changes in the host, thereby allowing resistance to persist even in the absence of the original plasmid. Considering this, the presence of the *erm*(55)^c^ gene on the chromosome of *M. chelonae* has been reported.^7^ Furthermore, in this study, the *erm*(55)^P^ gene was located within the HGT region and some plasmids carried genes predicted to encode products involved in genetic recombination, which may contribute to genomic rearrangement. Therefore, considering the potential role of plasmid– chromosome interactions in the long-term maintenance of macrolide resistance may be important.

This study had several limitations. The isolates were obtained from a single testing laboratory and, therefore, may not reflect nationwide trends in Japan. Although *rrl* point mutations associated with resistance in other *Mycobacterium* species were not detected in *M. murale*, resistance was observed on day 3. However, in the present *M. murale* isolate, the underlying mechanism remains unclear and requires further investigation. Despite these limitations, this study demonstrated the existence of macrolide-resistant RGMs related to the *erm*(55) gene-carrying plasmid in Japan. In the treatment of RGM infections, confirming inducible macrolide resistance is essential for accurate interpretation of MIC values, while plasmid-mediated resistance warrants further attention as an additional concern. Concerns have been raised regarding the spread of this plasmid and the *erm* genes among RGMs. Further investigations worldwide are warranted to thoroughly assess the spread and impact of this macrolide resistance-conferring plasmid.

## Transparency declarations

### Data availability

The complete chromosome and plasmid sequences for SRL2021-127, SRL2019-498, SRL2021-291, SRL2023-024, and SRL2023-035 were deposited in the DDBJ/ENA/GenBank databases under accession numbers AP043661 and LC872744, AP043662 and LC872745, AP043663 and LC872746, AP043664 and LC872747, and AP043665 and LC872748, respectively.

## Acknowledgments

We would like to thank Akiko Yamashita, Yukari Nogi, and Ginko Kaneda (National Institute of Infectious Diseases) for their assistance, and Editage (www.editage.com) for English language editing.

