## Supplementary methods for "Plasmid-mediated macrolide resistance among rapidly growing mycobacteria in Japan"

PCRs in this study were performed in a total of 20 µL mixtures containing 12 µL AmpliTaq Gold 360 with GC-enhancer (Thermo Fisher Scientific, Waltham, MA, United States), 6 µL $H_2O$, 0.5 µL of 10 nmol forward primer, 0.5 µL of 10 nmol reverse primer, and 1 µL of bacterial suspension as a template under PCR cycles: an initial denaturation step of 10 min at 95°C for the activation, followed by 40 cycles of 95°C for 40 seconds, 60°C for 40 seconds, and 72°C for 60 seconds with a final extension step of 72°C for 10 minutes. PCR products were visualised on an agarose gel by electrophoresis.

To identify species within several complexes, whole-genome sequencing analyses were performed. DNA extraction, short-read sequencing, and genome assembly were performed as previously described,[1] with sequencing conducted using the NovaSeq X Plus platform (Illumina, CA, USA). A taxonomy check of genome sequences was conducted using average nucleotide identity analysis with DFAST_QC v0.5.7.[2]

In the pangenome analysis, a Venn diagram was generated by calculating and drawing custom Venn diagrams (https://bioinformatics.psb.ugent.be/webtools/Venn/) (Figure S3). Core alignment from Roary v3.13.0 was trimmed using trimAl v1.4. rev15 with the option '-automated1'.[3] A maximum likelihood tree was constructed using the best-fitted nucleotide substitution model (HKY+F) in IQ-TREE v2.3.6,[4] with a 1,000-replicate, ultrafast bootstrap approximation.

Plasmid sequences were analysed using Snippy v4.6.0 (https://github.com/tseemann/snippy) and Gubbins v3.4[4] and a recombination-free phylogeny was generated in IQ-TREE v2.3.6, with the option '-m GTR+ASC' and a 1,000-replicate, ultra-fast bootstrap approximation. Phylogenies were visualised using an Interactive Tree of Life[5] (Figure S3).

### Supplementary figures

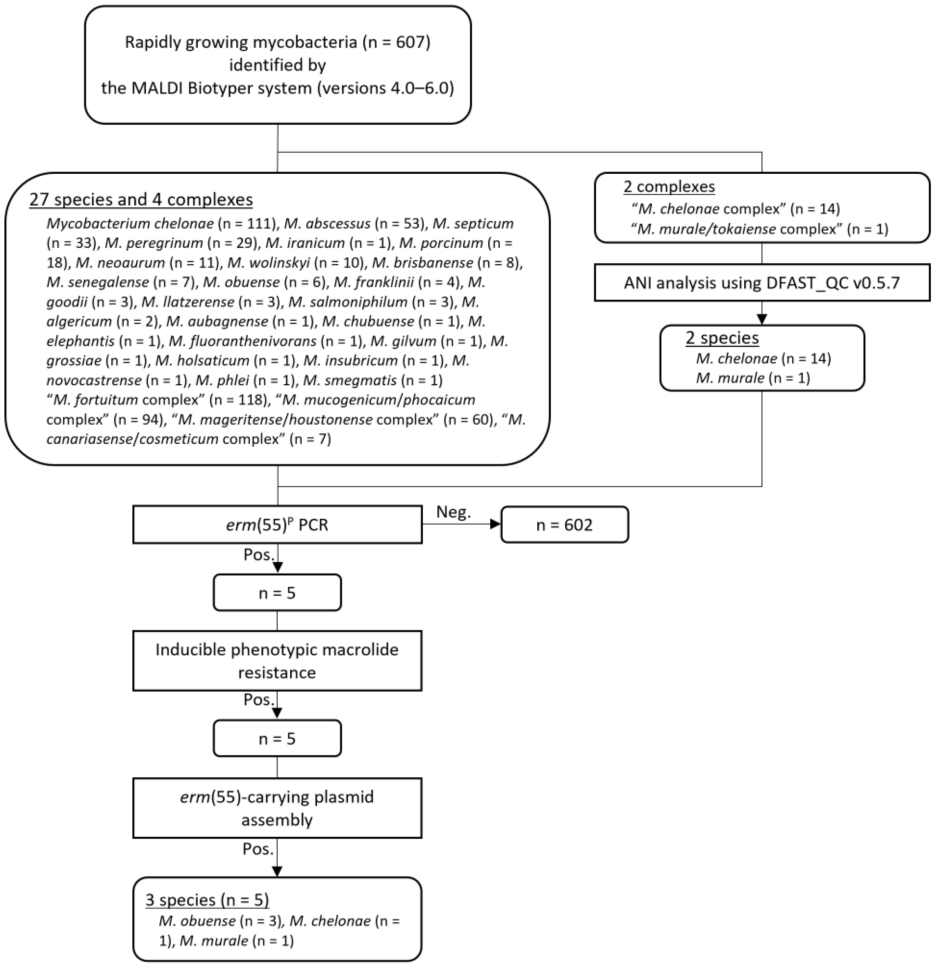


Figure S1. Flowchart of the screening process for the plasmid containing the *erm*(55) gene. ANI, average nucleotide identity.

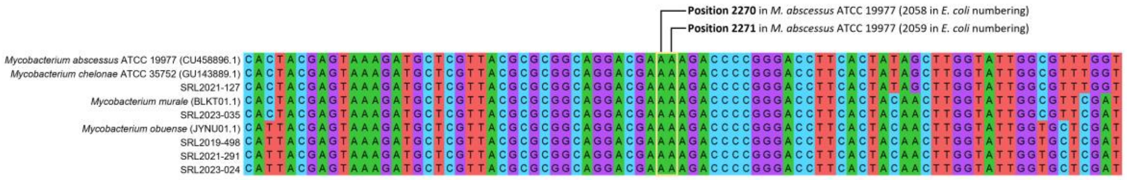


Figure S2. Alignment of 23S rRNA (*rrl*) genes from *erm*(55)-positive isolates and reference *Mycobacterium* strains. Nucleotide positions 2058 and 2059 (*Escherichia coli* numbering), implicated in acquired macrolide resistance, are highlighted in yellow.

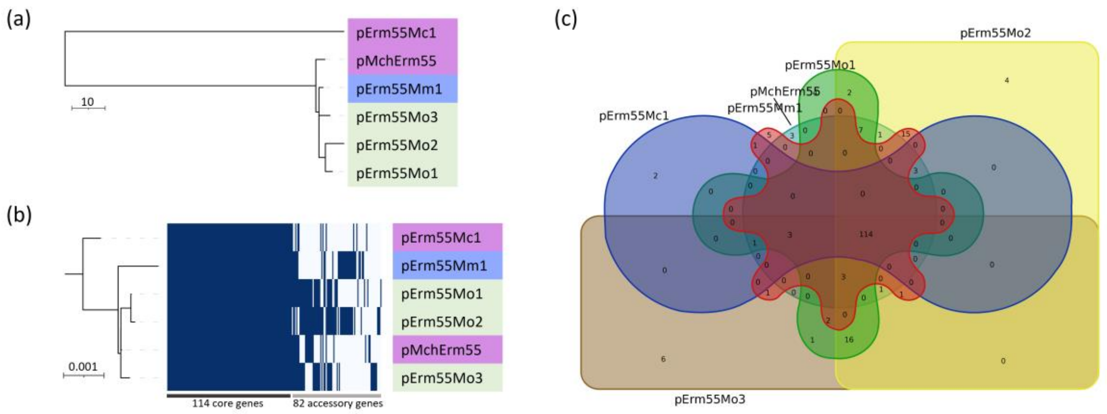


Figure S3. Comparison of *erm*(55)-carrying plasmids. (a) Recombination-free phylogeny. The scale bar indicates the number of SNPs. (b) Phylogeny based on core gene alignment. The scale bar indicates the number of substitutions per site. (c) Venn diagram of core and accessory genes. Among the 196 genes identified, 114 were core genes shared between pMchErm55 and plasmids carrying the *erm*(55) gene assembled in this study.
